# System Environmental Metrics Collector for EM facilities

**DOI:** 10.1101/2021.11.04.467268

**Authors:** Lambertus Michael Alink, Edward T. Eng, Robert Gheorghita, William Rice, Anchi Cheng, Bridget Carragher, Clinton S. Potter

## Abstract

Recent developments in cryo-electron microscopy (cryoEM) have led to the routine determination of structures at near atomic resolution and greatly increased the number of biomedical researchers wanting access to high-end cryoEM instrumentation. The high costs and long wait times for gaining access encourages facilities to maximize instrument uptime for data collection. To support these goals, we developed a System Environmental Metrics Collector for facilities (*SEMCf*) that serves as a laboratory performance and management tool. *SEMCf* consists of an architecture of automated and robust sensors that track, organize and report key facility metrics. The individual sensors are connected to Raspberry Pi (RPi) single board computers installed in close proximity to the input metrics being measured. The system is controlled by a central server that may be installed on a RPi or existing microscope support PC. Tracking the system and the environment provides feedback of imminent issues, suggestions for interventions that are needed to optimize data production, and indications as to when preventative maintenance should be scheduled. The sensor components are relatively inexpensive and widely commercially available, and the open-source design and software enables straightforward implementation, customization, and optimization by any facility that would benefit from real time environmental monitoring and reporting.

## Introduction

Producing high-resolution cryo-electron microscopy (cryoEM) reconstructions of bio-macromolecular complexes often requires acquiring thousands of images. Despite recent progress in automation and throughput (Baldwin, Tan et al. 2018, Cheng, Eng et al. 2018, Eisenstein, Danev et al. 2019, Lyumkis 2019) access to the highest end instruments remains scarce. Any reduction in throughput capabilities adds delay to experimental timelines. This situation emphasizes the need for workflows that maximize experimental throughput and are verified by routine benchmarking of the microscope performance (Kim, Rice et al. 2018). Currently, the accepted practices for microscope standard operating procedures vary significantly between facilities. Whether in an academic or industrial setting each site has its own strategy aimed at optimizing the performance and the uptime of the instruments (Alewijnse, Ashton et al. 2017, Sader, Matadeen et al. 2020). One common theme that all these strategies share is to create an operational environment where instrument stability is rigorously maintained over the course of a data acquisition session.

High-end microscope hardware is currently capable of minimizing imaging aberrations and distortions in order to ensure the highest spatial resolution (Rose, Nejati et al. 2019). However, the system is still dependent on several ancillary components including consumables (liquid nitrogen), support equipment (computers), building infrastructure (air handlers, chillers, and air compressors), and the ambient environment (temperature, humidity, vibration, electro-magnetic fields) that contribute to operational variations. Monitoring the performance metrics of these components can be used to extract useful information and gain insight into the daily operation of the high value instruments in the facility. This feedback enables instrument performance to be optimized and throughput to be maximized.

When creating an environment that allows high-end instrumentation to operate stably at a high level of throughput it is important to understand that systems vary in their maximum requirements. The higher the resolution required from an experiment the greater the microscope’s sensitivity to the room it is sited in, including magnetic, mechanical, acoustical and thermal disturbances (Muller, Kirkland et al. 2006). Prior to the installation of high-end equipment, vendors will conduct a survey on the thermal, electromagnetic and vibrational environment to determine the nominal conditions at the site location and if these allow for optimal instrument performance. During room construction there are best practices that will mitigate environmental instability (O’Keefe, Turner et al. 2004). However, even if the room environment and stability exceed the vendor requirements there will still be daily variations.

Here, we present the design of a Systematic Environmental Metrics Collector for facilities (*SEMCf*) and demonstrate how continuous monitoring and reporting of metrics, including liquid nitrogen (LN2) tank levels, room temperature and humidity, chiller temperature and pH, and EMI cancellation systems, have contributed to improving the reliability and efficiency of the instrumentation housed at the New York Structural Biology Center, which currently supports twelve transmission electron microscopes. SEMCf utilizes the well-known and economical Raspberry Pi (RPi) platform. This device allows dedicated sensors and processing software to be installed close to the input metrics being measured. For facility staff, online web pages provide instant snapshots of the current status and various reporting tools that can be used to understand barriers to efficiency. In the case of an outlier being detected in the facility support infrastructure that might cause sample loss, the software automatically sends an instant message via Slack^1^ to alert the staff.

The system construction is straightforward using standard commercially available components and controlled by Python programs run on a central server. As described, the system is scalable to monitor multiple instruments in a large and complex facility environment. The designs for all of the components are freely available as open source files. This design philosophy should enable other research groups to easily adapt the system to collect a variety of metrics to optimize performance according to the needs of their specific facilities.

## Materials and Methods

The *SEMCf* system is comprised of three main hardware components: Raspberry Pi (RPi) single board computers, sensor units, and a central control server (Figure 1). The RPi single board computer (raspberrypi.org) runs the default Raspbian operating system and custom designed Python software to acquire data from the sensor units. The smart home sensor kits for the RPi system come standard with sixteen units that can monitor a variety of environmental metrics including temperature and humidity, sound, light, smoke, motion, flame, vibration, air pressure, CO level and water. In addition, adaptor kits allow connection with readily available laboratory sensors^2^ such as a pH probe to monitor microscope chillers or weight scales to measure LN2 tank weights. For each transmission electron microscope equipped with an autoloader, the base *SEMCf* deployment uses local sensor units monitoring the electromagnetic fields, room temperature, relative humidity and the LN2 tank weight. The LN2 supply tanks feeding the microscopes are placed on top of a RW-1000 scale (Cardinal/Detecto, Webb City, MO), which is capable of sending weight information through a serial cable. From the RPi the collected data is cached locally, then sent to a queryable database on a central storage server. All connections are hardwired to a local intranet to maintain security and minimize potential disruptions from WiFi network connection outages.

**Figure 1:**
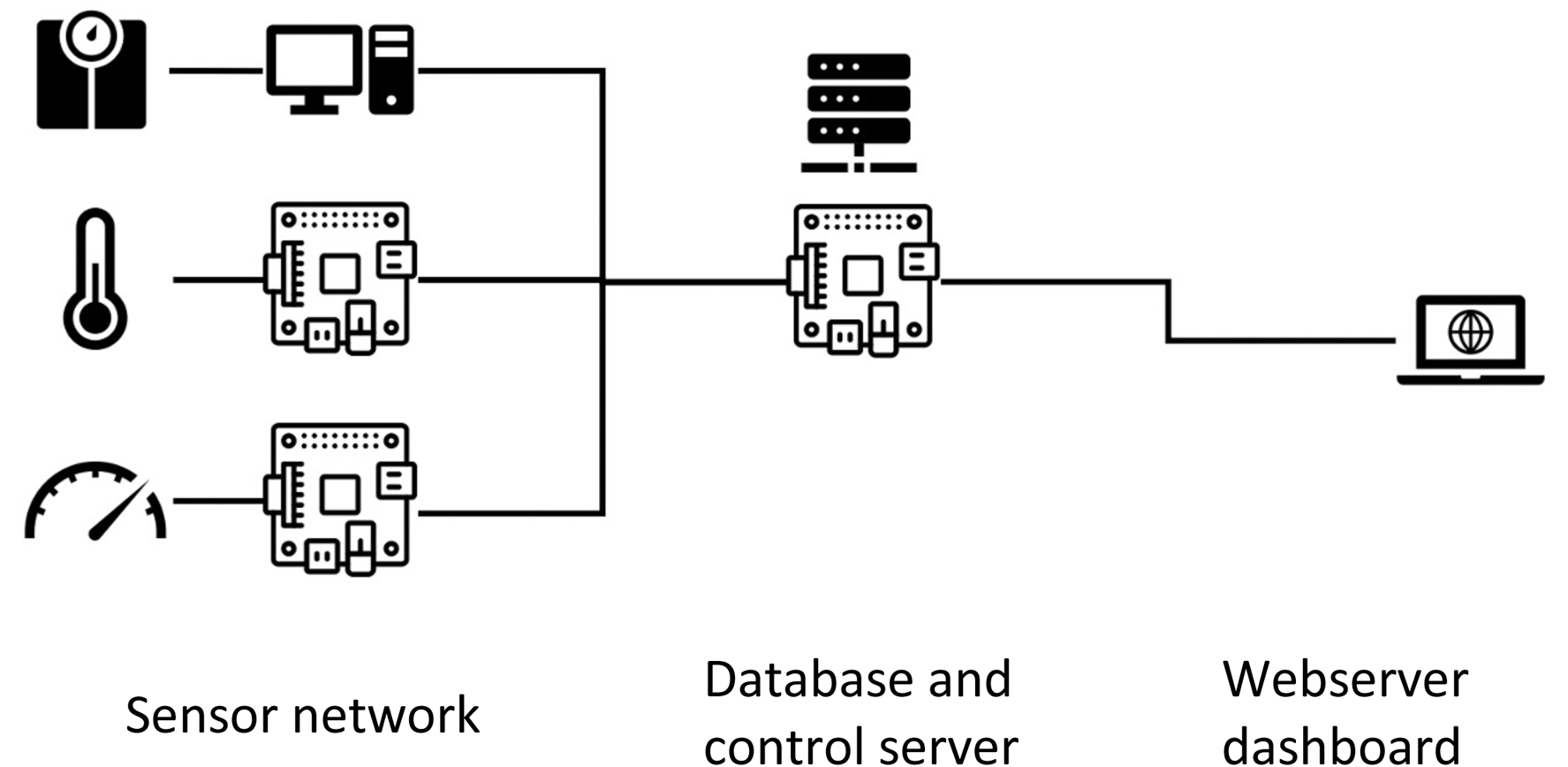
Architecture of the SEMC facility monitor. The network is built around RPi computers that connect to dedicated sensors for data collection. The data is fed to a central database running on a control server managed by Python control software. The network is accessible through a webpage allowing end user control and data visualization. User notifications are sent through email and Slack. All components are connected over wired intranet to the web server. VPN allows access to the networked RPis from outside the facility. The software is multi-threaded to be responsive to network requests and data handling.

The control software is written in Python and is compatible with Windows, Mac OS, Linux and RPi platforms. The primary control is divided into software packages for the environmental metrics clients (sensor package) and the master control server (control package). The *SEMCf* code repository may be found on: https://github.com/nysbc.

### Sensor packages

The RPi devices are used to manage the I/O for the ambient sensors. The RPi runs a Debian Linux operating system. At program startup, the Raspberry Pi loads the drivers for sensors and does an integrity check by confirming that a connection to the sensor can be established and data is received. If no sensor is found, the program will retry to connect to the sensor and stop after a timeout of 5 minutes. The RPi also establishes a connection to the central server. Upon receiving sensor data, a date and time stamp and device IP are added for logging and troubleshooting and the data is uploaded to the server. The temperature and humidity sensors are RPi home sensor modules connected to the RPi by pull-up resistors through a general-purpose input/output (GPIO) board that comes standard with an RPi kit. The LN2 supply tank weight scales are connected to the Thermo Fisher Scientific (TFS) microscope support PC that is delivered with a Titan Krios. The PC runs under windows and its serial 232 port is readily available to connect to the weight scale. The RPi may also be used by attaching a USB to serial port converter. On the support computer, the pyserial software library is added to the Python package to provide the interface for communication to the weight scales over the serial port. The python software package runs an integrity check on collected data and establishes ethernet connections to the central server. The sensors for temperature, humidity, water pH, and facility voltage, utilize the same client architecture, though using their respective specific software library to provide needed interface algorithms.

### Control package

The central server currently runs on a RPi but has also been tested on a Windows and Mac OS platform. The control package keeps track of active sensors and monitors if active connections have updated sensor data available. It checks data for alarm events or thresholds and sends warning emails once triggered. It also stores an archival database and hosts the web interface that provides access to historic data. All three functions are within the control package installation but are designed for multi-threading to be responsive to new network requests while handling data in parallel. Critical software parts are programmed using exception capture and auto-restart. The Python code checks connection quality between sensors and central server and if a sensor is no longer available it is deleted from the list and no longer polled for updates.

## Results and Discussion

### Communication and database metrics

The System Environment Metrics Collector for facilities (*SEMCf*) is currently deployed at two sites: the Simons Electron Microscopy Center at the New York Structural Biology Center (NYSBC) and the New York University Langone Health Cryo-Electron Microscopy Laboratory (NYU Langone). The system is configured to monitor the facilities to support eight transmission electron microscopes (TEMs) at NYSBC (seven Titan Krios and one Glacios) and two TEMs at NYU Langone (one Titan Krios and one Talos Arctica). The system monitors the LN2 supply of the microscopes by sending alerts in case of tank failure, frozen depressurization valves that lead to greater LN2 consumption, and warnings before a tank runs empty. The aim is to prevent cryo sample loss and damage to system vacuum components due to the system warming up from lack of LN2. At NYSBC, in addition to LN2 supply tank monitoring, the system is designed to monitor humidity, ambient temperature, and the pH of the chilled water systems. These added features were helpful as NYSBC recently underwent an upgrade and expansion. When additional microscope rooms and facility environmental controls were added, it changed the HVAC balancing of the microscope suites and these could be understood from the *SEMCf* provided feedback of absolute room temperature and maximum temperature swings per 24 hours.

*SEMCf* is a modular platform with a sensor network and control system (Figure 1). Sensor deployment is arranged in such a way that control and readout of environmental factors is conducted as close as possible to the monitored microscopes. By installing RPi computers in the microscope room, the environmental monitoring network can easily be expanded with additional sensors without the need for extensive rewiring. Each RPi computer can be monitored remotely over VNC and software can be updated if needed. The system will attempt to self-heal by remotely power cycling sensors in error states. Also, if there is a power failure or interruption, the sensor monitor software will automatically reboot and attempt to reconnect to the central server. However, if one sensor is unresponsive it does not affect the ability of the sensor network to report metrics. In the case there is a failure of the central server, the system will temporarily stop reading sensor information. Once the server is online, it recovers its last known good state from the database and attempts to recreate the network to the sensors to restart monitoring of the facility.

The database and control server provides real time alerts and longer term visualization of trends (Figure 2). Operationally, the RPi constantly cycles through all sensors that are online. Once any of the sensors shows an outlier, such as high room temperature or LN2 consumption out of expected range, the server automatically generates an event. The threshold to turn this event into an alert may be set in various ways, for example as a manually defined allowable temperature window or as an unexpected rate of LN2 use. The LN2 monitoring system has a lookup table drawing from a historical database accumulated during more than a year of monitoring LN2 supply tank use for six Titan Krios instruments at NYSBC. If the event triggers an alert, then email and Slack notifications are sent to the operational staff to allow interventions to mitigate possible damage or grid loss due to environment failure. In addition, the control server also hosts a web page graphical user interface (GUI) displaying a graphical dashboard of all facility metrics (Figure 2). The GUI also allows facility staff to set customizable parameters like tank tare weights and alert thresholds. The backend of this web site is the control software package that presents a manually defined rolling window of recent metric data and automatically filters out apparent outliers, such as negative relative humidity values or other spurious readings from the environmental sensors.

**Figure 2:**
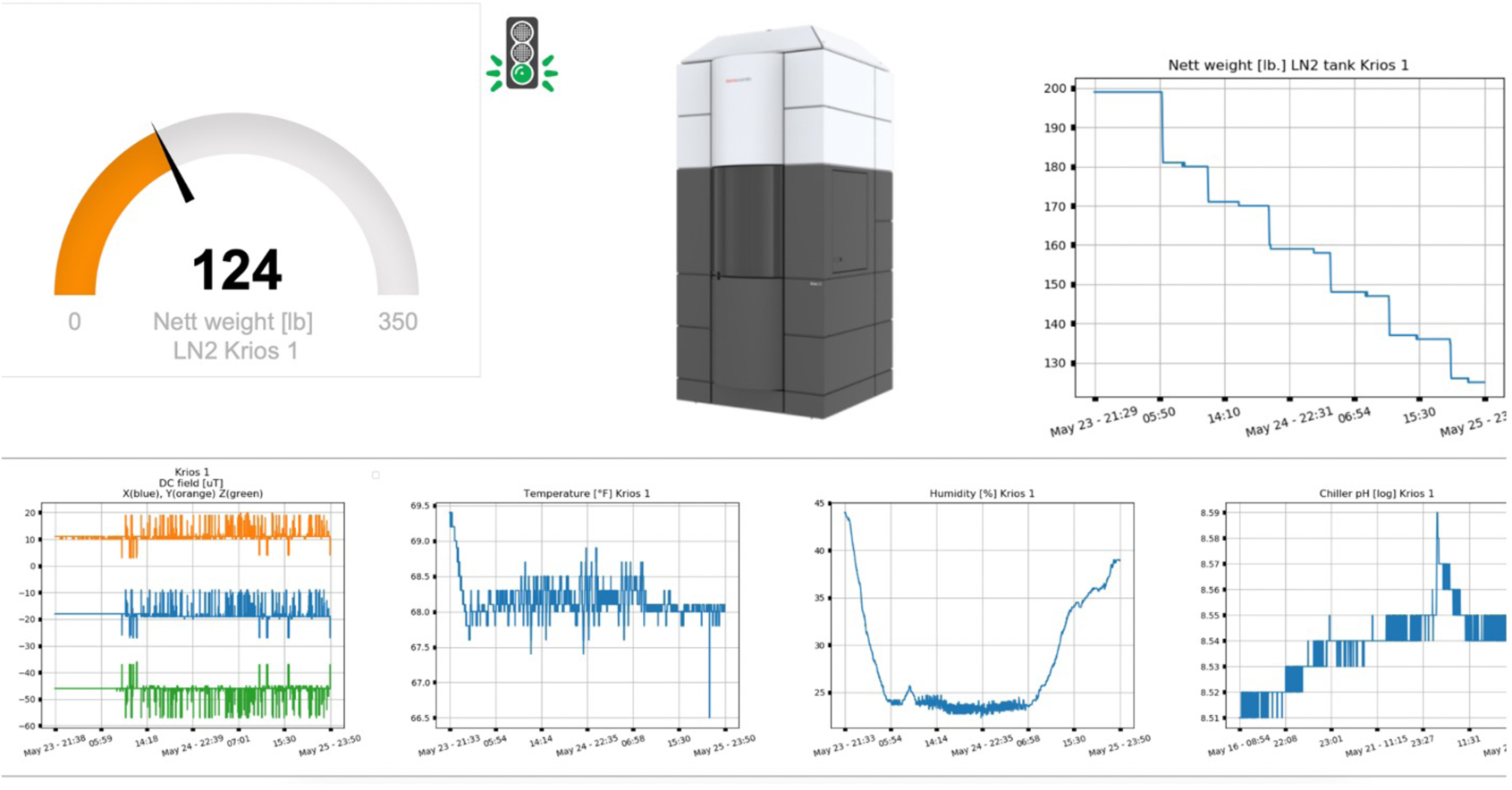
The central server allows real time monitoring. Alerts can be set up and a web page is hosted to display critical information about the facility. The control interface also allows more accurate monitoring by allowing for editing various parameters like the LN2 tank tare weights and error limits for alerts. The database can display data from any user selected timeframe.

The system specific data should be compared against data from all systems in the installed base to identify outliers that may be specific to the locale. For example, at NYSBC there are two HVAC air handlers, a primary and a secondary unit, feeding a group of microscope suites to maintain environmental stability. When there is a cycle from the primary to secondary unit for load balancing there is a temporary minor fluctuation (a spike in temperature of less than 1°F over 15 minutes) as one unit ramps down while the other ramps up. More specifically, because of the various air duct runs, certain rooms experience this event earlier than other rooms and have less of a temperature fluctuation. This event in its totality can be caught automatically and programmed to not trigger or send an alert during this process. One value of this database is to correlate deviations in cryoEM datasets (high drift or energy filter slit instability) at some point in time to room temperature outliers so as to aid in the analysis of failures. Overall, brief changes in temperature are considered less harmful, and are ameliorated by the instrument enclosure, than longer term drifts in temperature which may indicate hardware failure. A summary report of the current status, and any outlier events, can be configured to be sent out by email or Slack on a set interval (e.g. beginning of each day) (Figure 3).

**Figure 3:**
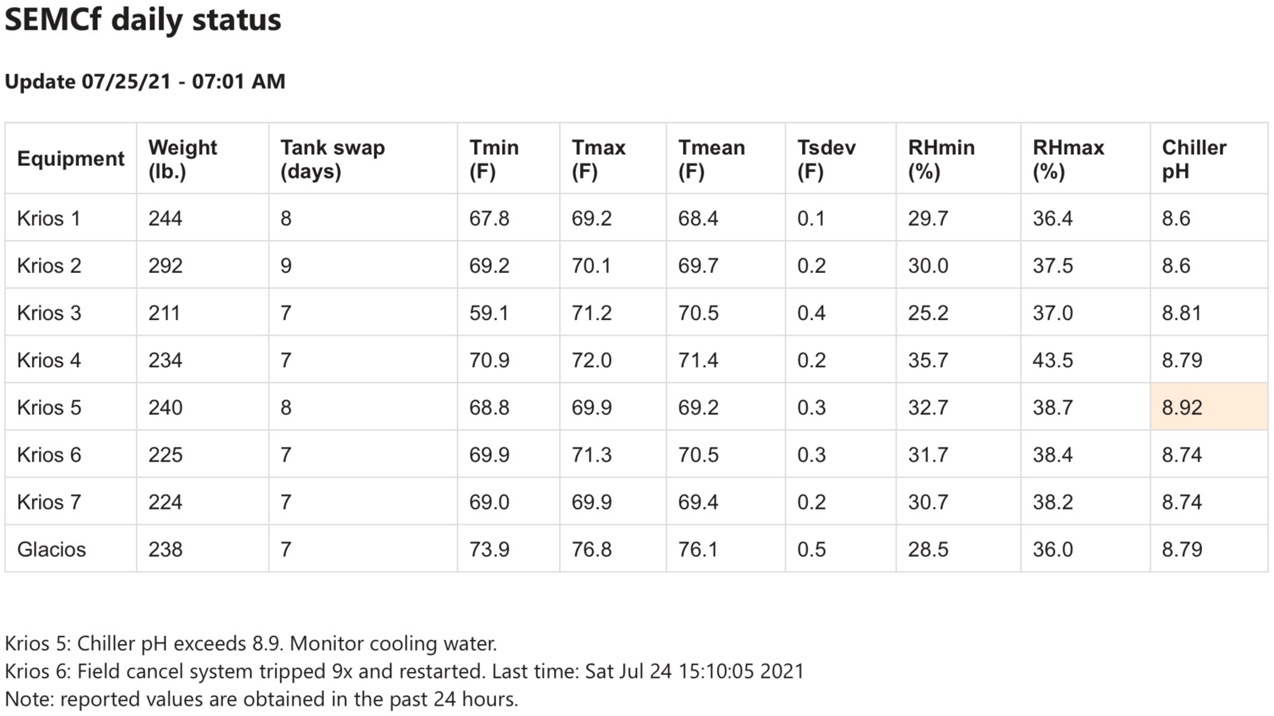
Daily emails provide a SEMCf overview of sensors and outliers.

### Liquid nitrogen cycle for microscopes

For facilities to maintain cryoEM operations, a critical consumable that needs to be tracked is LN2, which is used to maintain cryogenic (cryo) temperatures. High-end equipment, such as the Thermo Fisher Titan Krios and Glacios and the JEOL CRYO-ARM, are usually directly connected to an LN2 supply tank that the instrument uses for automatic filling of the microscope’s column and autoloader/sample loader dewar. After a certain period of time the system needs to be warmed up (conditioning cycle or cryo-cycle) to expel water that has accumulated over time on cold surfaces within the microscope column. To maximize throughput and minimize costs the goal is to minimize the frequency and duration of these conditioning cycles and use each LN2 supply dewar to its full potential while providing an adequate safety margin.

Facilities use various strategies for supplying LN2 to an instrument. For example, facilities may have bulk tank reservoirs, or a manifold that combines several LN2 supply tanks together. A standard configuration recommended by vendors in their instrument pre-install manuals is to use a single 160/180L or 230/240L LN2 supply tank. The exact tank configuration may depend on the gas supplier and institutional operational health and safety regulations. The desired volume of these tanks is to allow operations to last a week but may vary due to the amount of LN2 required or the integrity of the supply tank itself. In addition, if a cryo-cycle is scheduled, then there should be no LN2 fill ahead of it as it would waste product and extend the cryo-cycle time as the excess LN2 has to be boiled off. Conversely, if there still is LN2 in the system, then data collection may be extended, and the cryo-cycle delayed until the LN2 level naturally reaches the trip level for a fill. The product cost for a full rental LN2 supply tank from a gas supplier is relatively inexpensive (~ $75 in New York City). The main operational considerations are staff time required to change tanks, impact on data collection, and scheduling interventions within normal operating hours. At large facilities with over half a dozen autoloader systems, the cumulative tasks translate to significant overhead.

To execute a successful LN2 operational strategy using single tanks, real time weight monitoring is an important component (Figure 4). With this information the consumption rate may be monitored and used to troubleshoot systems if there are any deviations. The LN2 supply tank content is a combination of LN2 and gaseous N2. However, the LN2 density at 22 psi is more than two orders of magnitude higher than gaseous N2 and thus the measured weight is primary LN2. To capture this information the LN2 supply tank weights are measured using an electronic scale in one-minute intervals. This time interval provides sufficiently accurate statistics to track LN2 use of the normal operations of a microscope. A delivered 22 psi, 240 L LN2 supply tank has a tare weight between 300-360 lbs (135-165 kg) that must be accounted for to obtain an accurate measurement of the amount of LN2 product available. NYSBC uses rented tanks from a local supplier that are reused and may be held at the vendor for a period of time before delivery. The liquid valve of the LN2 tank is connected to the autofill system of the Titan while the gas valve is used to vent the autoloader to allow exchange of cassettes with samples into the autoloader. Note that the autoloader remains at LN2 temperature while loading a cassette, the gaseous atmosphere created by venting with N2 prevents excessive ice built up during docking.

**Figure 4:**
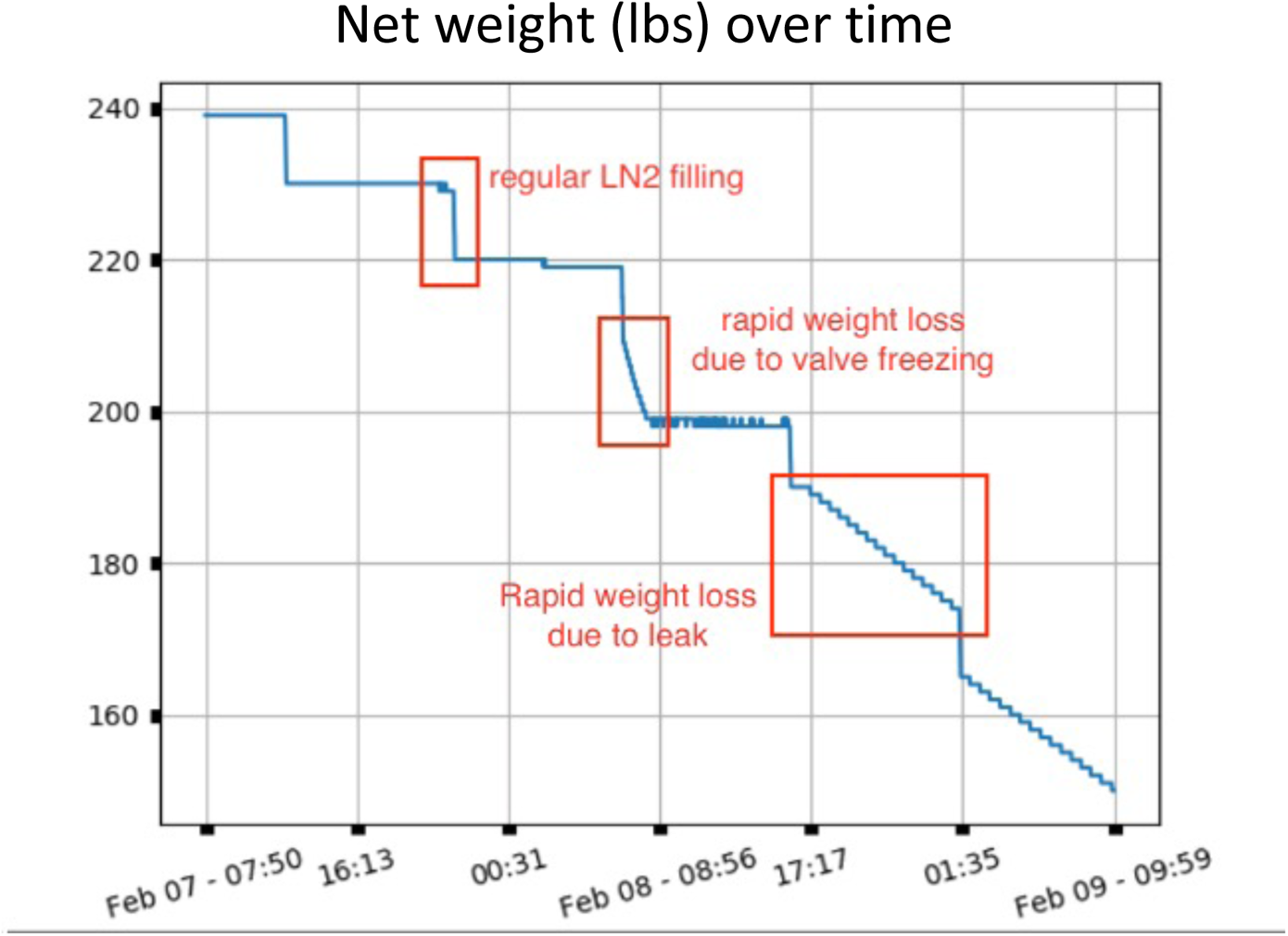
Continuous monitoring of LN2 tank weight provides insight into regular operations and failure events.

The pressure of the LN2 tanks is maintained by a mechanical bleed off valve to prevent over-pressure. Occasionally these pressure valves vent excessively and cause a drop of pressure that may impact the LN2 fill or autoloader sample exchange. Because of this, the LN2 supply tanks used at NYSBC also have a pressure builder that is used to raise the pressure back up to an acceptable level. For a 22-psi pressure tank there is a median product loss of approximately 5 lbs per day from the vent to maintain the pressure, but that may vary in a range of 3-7 lbs depending on the integrity of the tank (Table 1). An increased tank failure rate is observed when the tank is nearing empty (i.e. less than 75 lbs). With continuous weight monitoring the staff can therefore set this value as a lower trip level. In emergency situations, real-time monitoring allows systems to use tanks with as little as 10-20 lbs. of product remaining in the tank. The online feature again allows this to be monitored offsite after hours.

**Table 1:** **Average environmental conditions for the SEMC facility (November 2020)**

|  | **Average** | **Median** | **Max** | **Min** |
| --- | --- | --- | --- | --- |
| **Daily LN2 loss per day (idle state)** | 3.5±0.5 lbs | 5 lbs | 7 lbs | 3 lbs |
| **Daily LN2 use per day (maintain LN2 temperatures)** | 20±3 lbs | 23 lbs | 30 lbs | 16 lbs |
| **Daily temperature (facility wide)** | 69.7.±1.0 °F | 69.9 °F | 73.9 °F | 65.9 °F |
| **Daily relative humidity (facility wide)** | 22.5±1.0 %RH | 26.5 %RH | 48.5 %RH | 10.5%RH |
| **Daily temperature (best room)** | 68.2±0.2 °F | 68.3 °F | 68.5 °F | 68.1 °F |
| **Daily relative humidity (best room)** | 21.4 %RH | 26.7 %RH | 40.7 %RH | 12.7%RH |
| **ZLP stability over a 2 day period** | 0±0.4 eV | 0 eV | 1 eV | -1 eV |

The median LN2 consumption to maintain all nitrogen temperature for a Krios system is ~28 lbs per day with a daily range from 19-37 lbs of product. The upper end of the range is due to higher consumption when cooling the system down after a cryo-cycle. The microscope system triggers an autofill of LN2 when a dewar becomes low. The column dewar LN2 use is relatively stable, but the autoloader dewar tends to have a varying draw rate because sample loading requires venting of the system with gaseous nitrogen during introduction of new cassettes. Stabilizing the autoloader temperature after sample loading requires additional LN2 cooling power. The sensor network monitors the LN2 tank weight to predict tank empty and failure conditions, send alerts and advise on preventative maintenance actions that might be needed in case of outliers. This monitoring helps avoid catastrophic events like a sample loss due to the system warming up and also helps determine if tanks are being filled more often than necessary, which would negatively impact data acquisition throughput.

An LN2 autofill is typically started every 8 – 10 hours and stops within 15 minutes. The start and stops of an autofill are noted by the rate of weight decrease; in between autofills the LN2 consumption is expected to be very limited. If there is an issue with the pressure regulator or a leak in the supply lines, then an unusually fast rate of product use in a fixed time period will be observed. The first failure mode (issue with pressure regulator) is usually observed just after a filling cycle. Due to the design of the LN2 tanks the tank pressure regulator cools down during an autofill cycle and on faulty tanks freezes in an open position. This causes excessive nitrogen blow off and will deplete the LN2 rapidly. During normal operations there will still be a background consumption of LN2 (i.e. not used to cool the microscope) since the tank is not completely isolated from its environment. In addition, the autofill infrastructure (e.g. filling hoses, system dewars, connections to LN2 switching valves) are also likely to have some leaks. If the background leak rate is above a specified number than the system will raise an alarm.

If an autofill event is completed and does not indicate an issue with tank pressure, then the system gives an estimate of the next fill cycle and a prediction of when the LN2 supply tank will be empty. When the weight approaches the tare weight of the tank the system can determine if it is at a level that may run empty and inform facility staff to put the column in conditioning to preserve LN2 to keep the autoloader cold, or warm up the system using a cryo-cycle.

### Temperature and humidity monitoring

A key factor in ensuring operational throughput is maintaining microscope optical stability. As recommended by the manufacturer, a Titan Krios microscope room ideally has no more than 1.8° F (1° C) change in temperature over the course of a day. This environmental constraint keeps beam drift, lens aberrations and zero-loss peak deviations on energy filtered systems within vendor specified operational limits. The ability to ensure an environment that has as small a temperature deviation as possible is primarily realized by the room design and physical plant operations. When there is a variation in temperature or an issue with the building HVAC most facilities do not have advanced warning systems to alert the microscopy staff. With active monitoring of humidity and temperature, early warning alerts prompt operators if they need to take action (Figure 5) to allow data collection to proceed without interruption or intervene to preserve precious samples.

**Figure 5:**
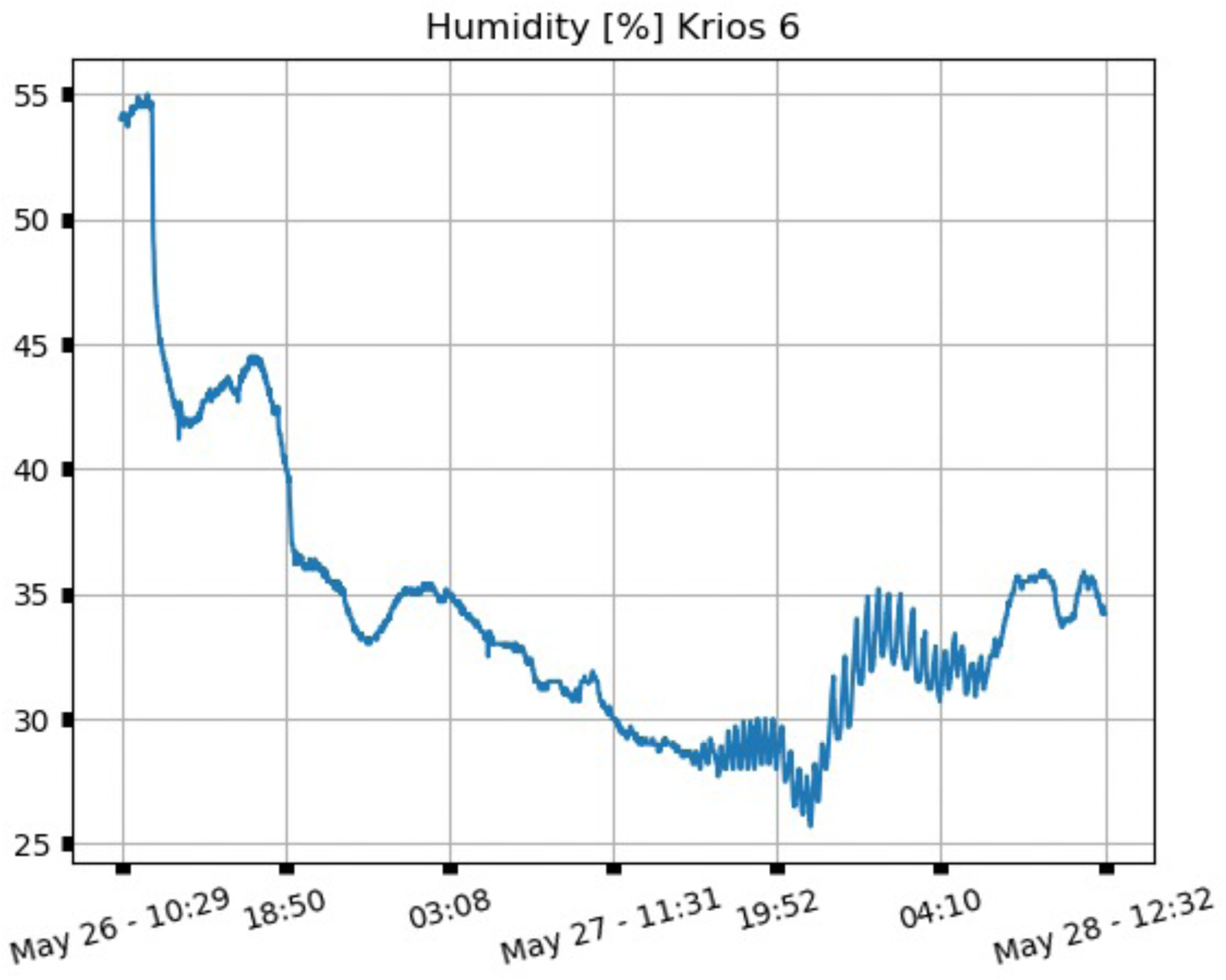
A faulty control board in the AC system caused the humidity to rise above 50% on May 26. These levels accelerated LN2 tank failure by freezing control valves. On May 27 the humidity control was restored, restoring active regulation.

SEMC’s autoloader microscope rooms are designed to be at a temperature of 68.8° F ± 1.2F across the facility with a daily maximum swing of 0.9° F (0.5° C) per room and a relative humidity (RH) within 7% RH swing at around 20% RH per room. However, the microscope rooms are not all the same size and are spread out across a facility with different HVAC systems which results in variations from room to room. The facility uses two different types of systems to maintain temperature and humidity. The older part of the center uses an HVAC system dependent on airflow, restricted in the microscope room by a duct sock, to maintain temperature and humidity. The newer rooms are supplemented by chilled beams, or radiant heating/cooling panels mounted on the walls of the room to control temperature as well as diffusers to circulate air through an inline desiccant wheel to bring down RH. While the instruments can tolerate a fairly wide range of humidity levels, lower humidity is an advantage when operating with cryogens and reduces condensation that develops over the course of the LN2 autofills.

One use case for *SEMCf* is due to weather conditions when the outside temperature changes by a significant amount (multiple degrees) within an hour. The HVAC system of the NYSBC facility is designed to maintain 20% RH. At 30% RH extra measures are engaged to maintain this humidity level. The *SEMCf* system tracks the variation in all the rooms and in failure conditions it will log a sharp rise in humidity and notify the staff. This is typically caused by a lag for the HVAC to switch on a dehumidifier in order to lower the humidity. Because different parts of the facility are controlled by multiple systems, in parts of the building the temperature can remain stable, while other parts show a temperature rise of 1° F in order to maintain 20% RH. The Krios enclosure will smooth out the temperature fluctuations and the *SEMCf* room monitor will send an alarm if there is a change of more than 1.6° F over 24 hours. Note that the change in room temperature often does not impact the microscope column dramatically as the beam drift is negligible, but the zero loss peak (ZLP) drift of the Gatan image filter (GIF), which is monitored through the logfile in Digital Micrograph, can still be found. If there are significant deviations in temperature, the operator can set the ZLP alignment to occur more frequently to prevent image loss due to occlusion by the GIF slit. Once the environment stabilizes, the operator can reset the interval between ZLP alignment checks back to nominal values. This ability to logically act based upon real-time room monitoring improves overall throughput and uptime. Some of these actions could also be automated.

### Electromagnetic field cancellation monitoring

To prevent electromagnetic fields from the environment compromising the resolution of the data, field cancellation systems are installed for all the Krios instruments at SEMC. The TMC Mag-NetX system is designed to create a magnetic compensation field by means of a current (max 3 A) in coil pairs (Helmholtz Coils) that are mounted along the room walls. The compensation field is equal and opposite to the disturbance field. The system uses a sensor installed in close proximity to the microscope column, inside the enclosure. The sensor is connected to a controller that together with the Helmholz coils creates an analog feedback loop. The controller aims to keep the error (the change in the signal) close to zero at the sensor location. The analog feedback loop allows for more precise real-time cancellation. Digital circuitry in the controller enables operational monitoring and remote control of the system via a USB or ethernet interface that allows the field cancellation system to communicate to and be controlled by the SEMCf system.

The SEMCf system monitors the Mag-Net-X system by taking samples every minute. The sampled data consist of the local DC field and slowly varying AC fields (<1 Hz). Typical values are around tens of micro-Teslas (uT) for the DC and few nano Teslas (nT) for the AC fields (Figure 6). Higher frequency AC EM-fields are typically blocked by mu-metal shielding in the enclosure and the Titan column.

**Figure 6:**
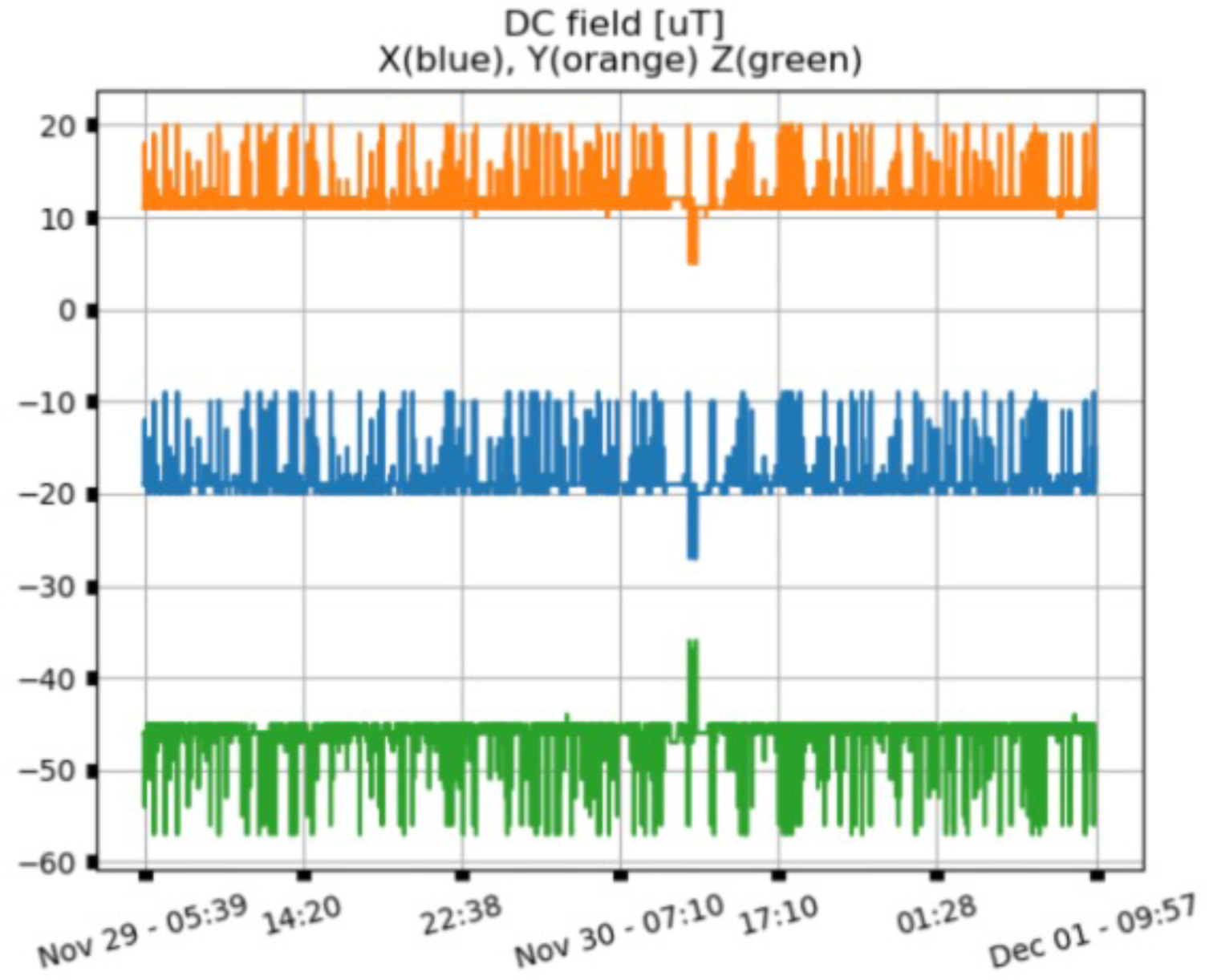
Mag-Net-X DC field measurement over two day period.

Remote monitoring allows checking if any saturation levels are reached (input fields on the sensor and output currents on the coils and alarm level on AC fields). Recently at NYSBC we had a use case where after servicing of the microscope the input antenna in the enclosure was displaced so that it was too close to the column. The signal picked up by the antenna saturated the controller about 10x per hour (>200 per 24 hour) during normal operation of the Krios. The supplier of the system was informed and after checkup the system was brought back to specifications and input signals calibrated. In more fatal error situations, an email is sent to the operators to attempt to restart the controller.

### Water loop and pH monitoring

To maintain a stable operating microscope with image movement (drift) below a specified threshold, the equipment’s temperature must be as stable as possible. For this purpose, each microscope has a dedicated chiller to control the temperature of the microscope and its environment. To properly specify the chiller it is important to account for the incoming and outgoing nominal heat flow to the instrument. A Titan Krios heat load depends on the specific configuration but may typically have up to two-thirds of its power (4000 W) dissipated into the microscope water loop whilst the remainder goes into the room around the microscope. The water-cooling system is thus a critical component for maintaining a stable room, microscope lens, and camera temperature. To mitigate small load changes the chiller has a two gallon reservoir that levels heat changes by absorbance due to the relative high heat capacity of water. Figure 7 provides an example where the chiller failed to maintain a stable temperature which may cause unacceptably high drift values of images taken by the affected microscope.

**Figure 7:**
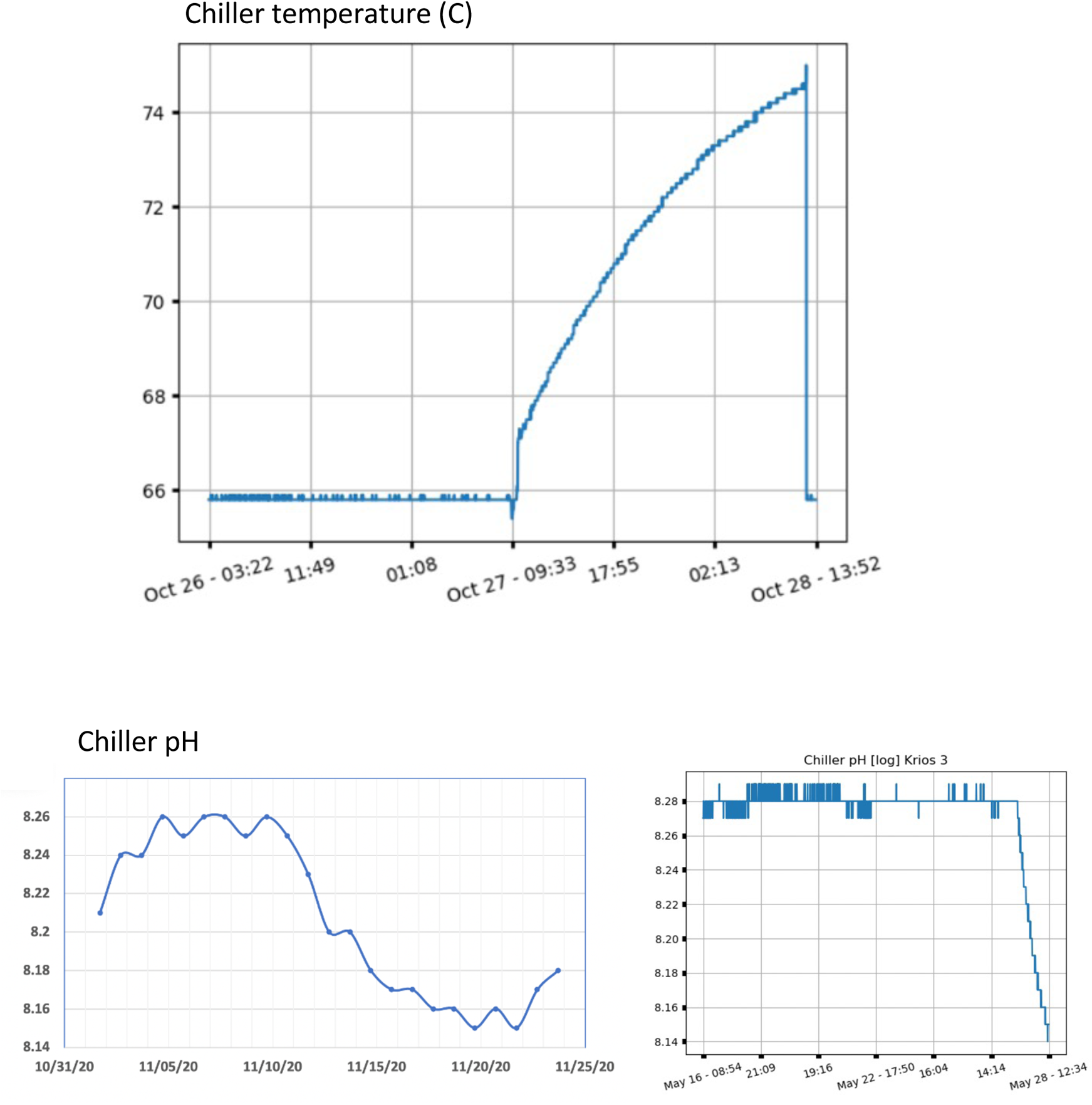
Top: Chiller failing to maintain stable cooling water temperature. Bottom: (left) Continuous monitoring of chiller pH proves early indications of water flow problems and (right) when insertion of a camera that had been idle for a few days causes changes in the cooling water recognized by the chemistry of the pH.

A dedicated RPi with a submersible temperature and pH sensor deployed in the chiller water reservoir allows continuous monitoring of temperature and pH level of the microscope’s water loop (Figure 7). The desired temperature and pH level of the system are recommended by Thermo Fisher Scientific (TFS). The recommended setting of the chiller temperature is such that the column temperature is midway between the room temperature and the water temperature. Typically the microscope is a few degrees warmer than its environment. The remaining heat that dissipates into the room is taken care of by the room air conditioning system.

The pH of the chiller water should be between 7.5 and 8.5 (ideal pH is 8.2) to prevent corrosion of the water system. During commissioning of a new water system or after replacing the water of an existing system there is a trend towards acidification - the pH is often buffered with NaHCO3, and over time, the removal of NaHCO3 by conversion to CO2 and outgassing causes a rise in pH. Typically, we measure a drop of 1 pH level each week for the first month, which then levels off. Changes greater than this amount may arise due to water pollution, biological growth, or some type of system degradation. Note that a low pH will allow more metals to dissolve into the cooling water. The *SEMCf* system flags changes in pH for investigation.

A major concern with a closed loop water system is algae, fungal and bacterial growth, which may be indicated by turbidity. The growth can degrade the accuracy of the flow gauges and limit cooling capacity. If left unattended, this contamination may affect the flow meters in the microscope water table by giving the appearance of flow in the normal range when in fact reduced or no water is flowing at all. As water in a circulation system ages it is exposed to air in the reservoir and absorbs carbon dioxide from the air making the water more acidic. This acidic water has an increased ability to extrude ions from plumbing systems and oxidize metal from pipes and fittings. This situation can lead to corrosion, for example green flakes (likely to be copper carbonates and/or sulfates). All of these issues emphasize that it is important to keep temperature and pH constant and as close as possible to the manufacturer’s limits to ensure stable operation of the microscope.

### Low temperature sample freezer

For sample storage, many facilities possess a low temperature sample freezer. Typically, these freezers store proteins at temperatures below −80 C. Samples are added by staff on arrival and removed just prior to grid preparation. Equipment failure or human error (e.g. improper closing of the door) can lead to inadvertent heating up of the freezer and loss of samples and thus a continuous monitoring and alarm system is needed. To monitor temperatures below −80 C, specialized sensors are needed since these low temperatures are inaccessible by off the shelf temperature sensors. For the Raspberry PI system, an interface is available that allows use of thermo couples to measure temperature. Virtually any type of thermocouple can be used in combination with the RPi, the interface board takes care of required temperature compensation and calibration. The RPi samples the data and makes it available for lookup, graphing and creating email or Slack alarms in case of malfunction.

## Conclusions

The *SEMCf* system provides a framework for daily reports and status updates on a variety of peripheral devices and environmental conditions in a facility. The core system can continuously monitor LN2 consumption, room temperature, humidity, stray fields, chiller water temperature and pH, freezer temperatures, room oxygen level and facility voltage. Alerts are sent if parameters are outside desired specifications. The RPi architecture allows for easy scalability and other capabilities can be readily added. This framework serves as an early warning system to address causes of loss of instrumentation throughput, data quality and downtime that would occur without preventative actions. Saving the metrics in a database enables facility staff to do an in-depth analysis of microscope output, support architecture, and environment of each instrument.

We have shown that a straightforward design combining inexpensive consumer components, is capable of impacting our ability to ensure availability of high-resolution cryoEM facilities through long term monitoring and resource management. All designs for the platform, parts, and the Python instrument control software, are available at https://github.com/nysbc. With these designs, scientists with limited access to technical workshops and equipment can easily construct their own system by making use of off the shelf commercial components. The intention of this open-source design is that the *SEMCf* system can be customized, modified and expanded for each specific facility’s applications. The remote accessibility of the central server and clients was a huge asset during the COVID-19 pandemic when onsite staff access was severely curtailed.

The *SEMCf* code repository may be found on: https://github.com/nysbc.

## Acknowledgements

Some of this work was performed at the Simons Electron Microscopy Center and National Resource for Automated Molecular Microscopy located at the New York Structural Biology Center, supported by grants from the Simons Foundation (SF349247) and the NIH National Institute of General Medical Sciences (GM103310) with additional support from Agouron Institute (F00316), NIH (OD019994) and NIH (RR029300).

## Footnotes

1 slack.com

2 CanaKit Raspberry Pi 4 4GB Starter Kit with Clear Case (4GB RAM) https://www.amazon.com/dp/B07YLY143F/ref=cm_sw_r_fm_apa_i_3Zs5DbA8S4XFX

